# Roles of Neuromedin-U and Endothelin-1 in human cardiac organoid differentiation

**DOI:** 10.64898/2026.10.23.761328

**Authors:** Rebecca R. Snabel, Carla Cofiño-Fabrés, Marijke Baltissen, Verena Schwach, Robert Passier, Gert Jan C. Veenstra

## Abstract

During recent years, many new culture methods have been established to generate cardiomyocytes, using signaling and patterning factors that play important roles in mammalian heart development. Here, we identify two additional pathways of interest using a cardiac differentiation roadmap that we have generated, involving the endothelin 1 receptor *EDNRA* gene, and the Neuromedin family signaling peptide gene *NMU*. Treatment with NMU had most effect in ventricular lineages where it normally is not abundantly expressed, leading to a significant shift from ventricular cardiomyocyte (vCM) differentiation to non-myocytes, as shown with reporters and cell type-specific markers, which was associated with an increased beating frequency. This was not observed in atrial cultures. Treating atrial cultures with endothelin 1 (ET-1, EDN1) increased the number of atrial cardiomyocytes (aCM), while reducing repolarization times, increasing the contraction frequencies and altering the patterns of action potential propagations across the tissue. In ventricular cultures, a reduction in ventricular cardiomyocyte (vCM) numbers was observed, suggesting a combinatorial effect of ET-1 and retinoic acid (RA) on cardiomyocyte differentiation. These results identify ET-1 and NMU as important signaling factors for cardiomyocyte differentiation and electrophysiology of cardiac organoids, with ET-1 enhancing aCM differentiation when added around the time of posteriorization of the second heart field. NMU added without RA reduces the number of vCMs in the culture while it changes the organoids to exhibit a more atrial-like beating frequency.

## Introduction

Cardiomyocytes (CMs) can be differentiated from pluripotent stem cells (PSCs) *in vitro*, as first established 40 years ago, using embryoid bodies (EB) obtained from mouse embryonic stem cells (Doetschman *et al*., 1985) and subsequently demonstrated using human PSCs (Itskovitz-Eldor *et al*., 2000; Zwi *et al*., 2009). A multitude of different protocols have been developed since, to culture CMs (Burridge *et al*., 2012; Mummery *et al*., 2012; Devalla and Passier, 2018). Most of these culture systems rely on signaling modulation to direct the cells towards cardiac lineages, and consist of a specifically timed addition of growth factors needed for cardiogenesis (Mummery *et al*., 2012), while also bringing along refinements such as a chemically defined and serum-free medium (Ng *et al*., 2008). During the initial steps of the EB culture, stem cells aggregate into spherical structures, in which mesoderm is induced and cells undergo an epithelial-mesenchymal transition (Chan *et al*., 2012; Zeevaert *et al*., 2020). Subsequently, cell aggregates are treated with BMP4, Activin A and the WNT-agonist CHIR99021, to induce precardiac mesoderm formation. During *in vivo* mesoderm development, cells are exposed to these factors at the primitive streak and by interaction with endoderm, a principle that was utilized by Mummery *et al*. in co-cultures with endodermal cells, to induce CM formation (Mummery *et al*., 2003). Furthermore, additional administration of retinoic acid (RA) to these cultures that contain precardiac mesoderm (at day 4), induces the posteriorized patterning of the cells, leading to a posterior second heart field (SHF) identity, and ultimately to the formation of CMs of the atrial subtype (aCM) (Devalla *et al*., 2015; Zawada *et al*., 2023).

Recently, we presented a cardiac roadmap of *in vitro* differentiation, spanning different culturing methods (Snabel *et al*., 2025). Within this map, we compared the EB cultures, or cardiac organoids, directed towards ventricular (non-RA) and posteriorized atrial (RA-treated) lineages. Following RA administration on day 4, the atrial and ventricular differentiation trajectories began to diverge. Within each lineage culture, additional pools of fibroblasts, endothelial and epicardial cells were identified and formed independent of the RA treatment and alongside the ventricular and atrial CMs (vCMs, aCMs). The direct comparison between the patterned cardiac organoids allowed us to identify CM lineage-specific gene regulatory networks, in which we could highlight the transcription factors responsible for regulating the cell type-specific transcriptomes. Upstream of these transcription factors, signal transduction cascades are presumably responsible for translating the signals coming from neighboring cells and other external factors present in its microenvironment. From day 4 onward, no external patterning factors are added to the cultures, relying on the acquired potency and cell-cell interactions for further differentiation. However, it is unknown what signaling cues cardiac progenitors need for efficient differentiation and what the source is of these signals.

In the current study, we aimed to define signaling factor differences between the atrial and ventricular cardiac organoid cultures, and to determine how perturbing specific signaling pathways affects their differentiation. We filtered differentially expressed genes across the lineages, for genes involved in signaling, and identified targetable factors of interest to use in perturbation experiments. Mapping these perturbation-induced transcriptional responses onto our *in vitro* differentiation roadmap allowed us to associate gene expression programs with specific cell populations and differentiation stages and to relate these changes to the observed functional phenotypes.

## Results

While the specification of cardiac progenitors has been investigated thoroughly, the lineage commitment of cardiac cell types is less well characterized. To analyze the potential for signal modulation in embryoid bodies, we re-analyzed the *in vitro* roadmap of cardiac differentiation that we developed recently (Snabel *et al*., 2025) (Figure 1A). This dataset comprised of 5,273 cells from multiple independent experiments (Figure 1B), spanning eight time points (day 4-21) and two culture conditions. In the absence of retinoic acid (RA), CMs in the cardiac embryoids adopt mainly a ventricular identity (vCM), whereas with RA treatment at days 4-5, aCMs are formed. We analyzed the time series data for differential signaling factors and their receptors, comparing the two lineages. We selected the clusters in each that contributed to the eventual CM pool and compared these for significant differences in gene expression (see Methods). These gene lists were filtered for signaling factors from the CellPhoneDB database (Efremova *et al*., 2020). The resulting compendium of ligand and receptor genes was visualized over time and per lineage and allowed us to define 59 genes grouped in 11 clusters with temporal expression patterns of interest (Figure 1C). Among these, gene clusters 11 and 7 represented early and late atrial signaling-associated genes, respectively, whereas clusters 1, 4 and 9 represented signaling-associated genes with more abundant ventricular expression.

**Figure 1:**
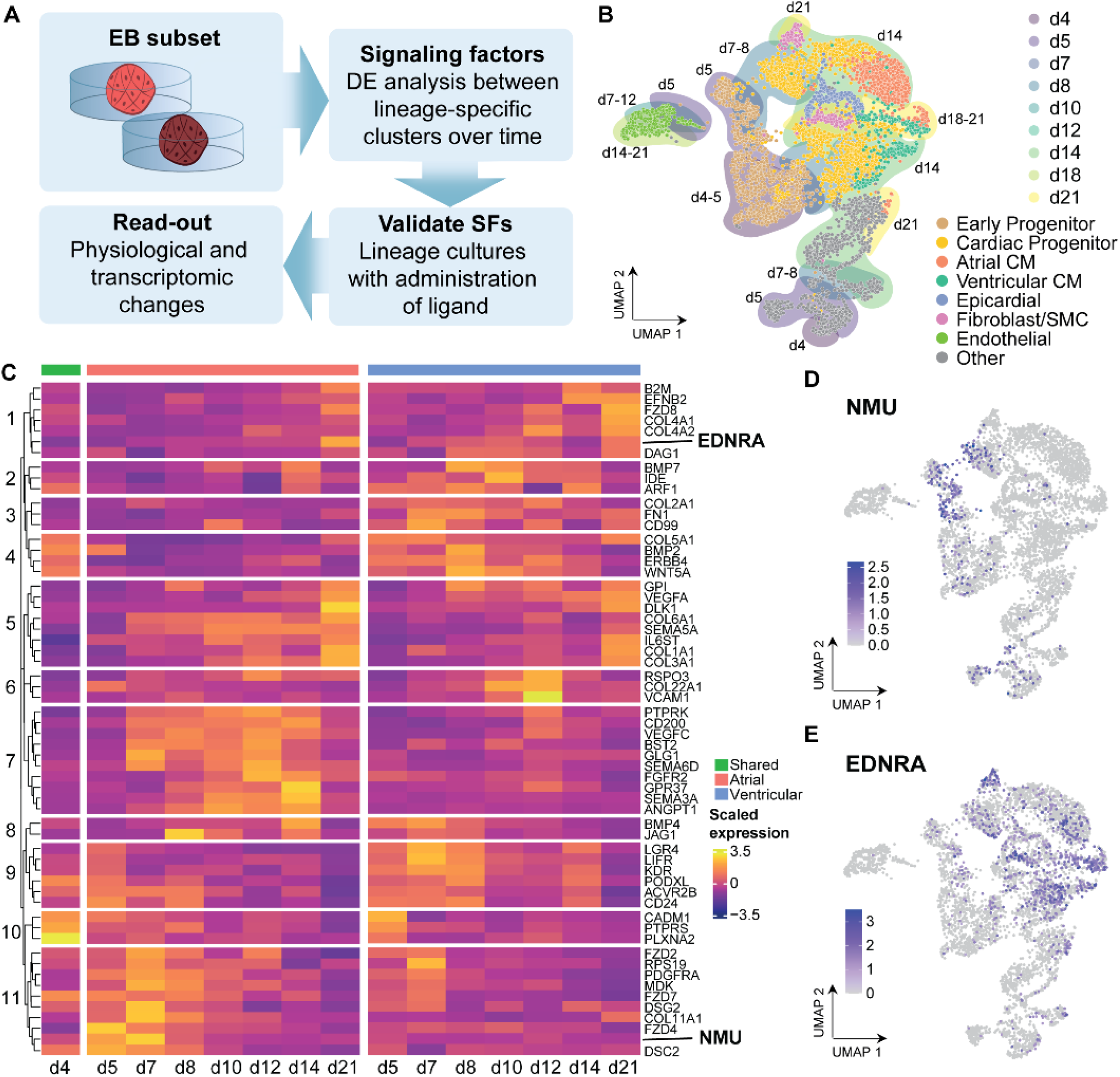
Signaling factor differences between the atrial and ventricular cardiomyocyte lineage. (A) Schematic overview of the analysis performed to select factors of interest. (B) UMAP with labelling of cell states within the EB subset of single-cell roadmap (Snabel et al. 2025). Color of the dots represent cells of the cell state in the legend, background color indicates day of differentiation. (C) Heatmap with differential signaling factors in atrial (pink) and ventricular (blue) EB cultures over time. (D-E) UMAP representation of the EB subset of the time-resolved single-cell roadmap (Snabel et al., 2025), with expression levels of the selected signaling factors of interest EDNRA (D) and NMU (E).

Further narrowing down of this list was performed by 1) manual exploration of expression patterns including that of the interaction partner of the ligand-receptor pairs, 2) identification of potential agonists or antagonists of the factors identified, and 3) checking the literature for a potential role in cardiac development or heart disease. Based on these criteria, we selected NMU and EDNRA for further validation experiments, for their atrial- and ventricular-specific expression patterns, respectively (Figure 1D-E). Not much is known about the roles of these genes in cardiac lineage-specificity, whereas they showed specific upregulation in either aCM or vCM cell clusters.

### NMU negatively impacts ventricular CM differentiation efficiency

The mRNA encoding the Neuromedin-U (NMU) neuropeptide was found to be specifically expressed in early atrial EB timepoints (day 5-7), alongside expression of its receptor *NMUR2* (Figure S1B). Although NMU expression has been detected in the heart, and NMU is known to be involved in regulating blood pressure and heart rate, among other functions (Tanida *et al*., 2009; Rahman, Shahid and Pilowsky, 2013), little is known about its role in cardiac differentiation. To study the role of NMU in lineage specification, we treated EB cultures with 10nM of NMU at day 4 of the atrial and ventricular differentiation protocols, mimicking the expression peak observed in the atrial lineage (Figure 1D). Treatment with NMU for 24, 48 or 72h from day 4 reduced the number of vCMs in the ventricular differentiation protocol (based on reporter expression, *NKX2-5*-eGFP-positive – *NR2F2 (COUP-TFII)-* mCherry-negative cells, Figure 2A-B), while it did not affect the aCM efficiency in the atrial protocol (*NKX2-5*-eGFP-positive – *NR2F2-*mCherry-positive cells, Figure 2C). This reduction in NKX2-5 expression resulted from a shift towards a population that was negative for NKX2-5 (from 9.7% in control versus 31% in NMU-treated vCM; Figure 2D-E). This effect was stronger after longer NMU treatment and was not observed under combined treatment with RA.

**Figure 2.**
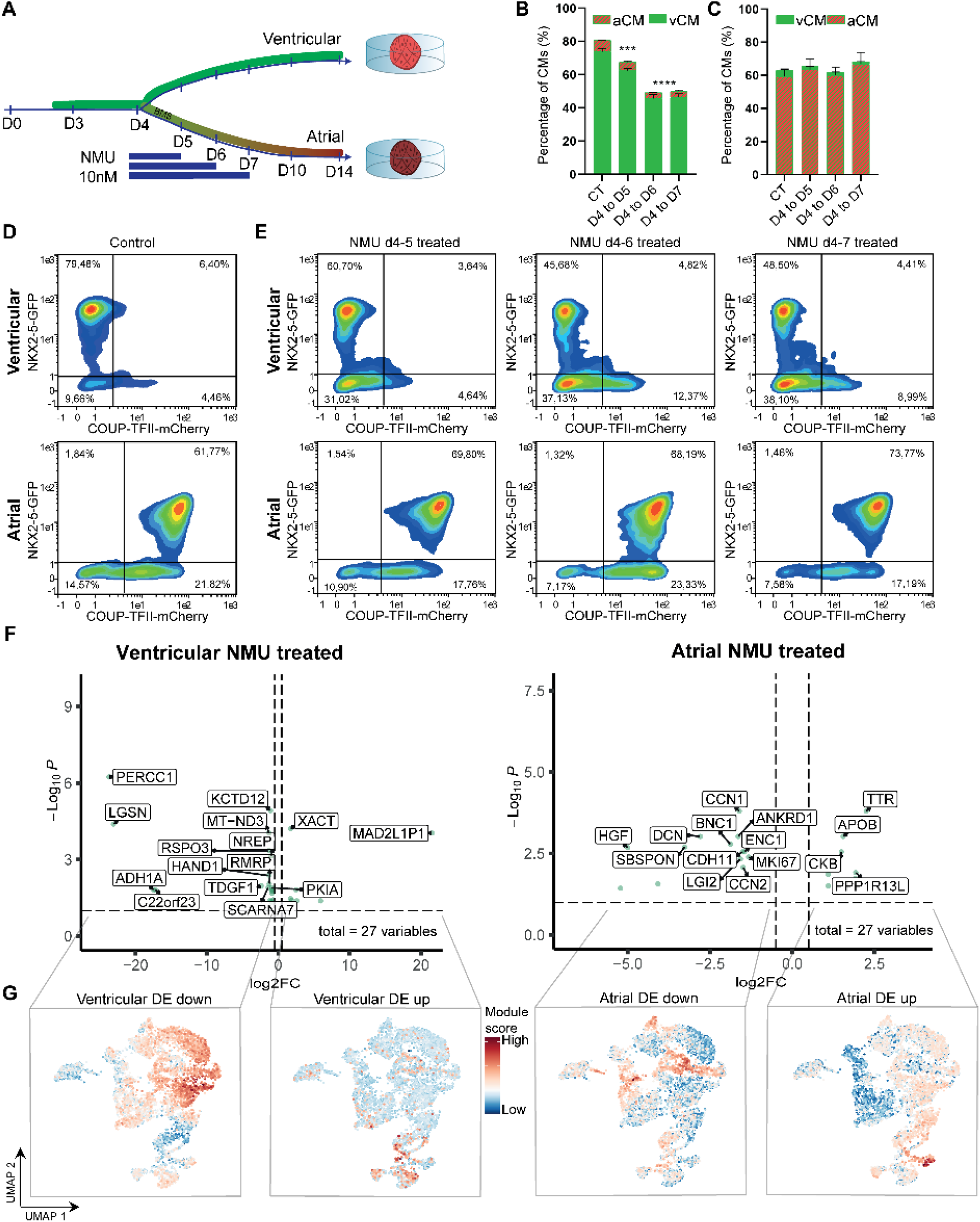
Effects of NMU treatment on cardiomyocyte lineages. (**A**) Schematic overview of lineage protocols with indication of different NMU treatments. (**B-C**) Percentage of reporters (aCM = NKX2-5- and NR2F2-positive, vCM = NKX2-5-positive and NR2F2-negative) as measured with FACS for the different NMU treatment timings versus no treatment (CT) in ventricular (B) and atrial (C) cultures; averages shown for triplicates. (**D-E**) FACS results with NKX2-5-GFP measurements on the y-axis and NR2F2-mCherry measurements on the x-axis as measured for the cells in the control situation (D) and under treatment with NMU for different durations (starting day 4 until day 5, day 6 or day 7, from left to right) (E). (**F**) Volcano plots representing the differentially expressed genes (p < 0.05) for the atrial (left) and ventricular (right) conditions treated with NMU for 24 hours. (**G**) The EB reference UMAP (Figure 1B) labeled with compound gene expression levels (as module scores) based on the lists of genes retrieved from (for plots left to right): non-RA down- (log2FC < 0), non-RA up- (log2FC > 0), +RA down-, and +RA upregulated genes, based on significance (p-adjusted < 0.05) upon NMU treatment.

To obtain a more detailed view of the effects of NMU treatment, we performed transcriptomic analysis (Figure 2F). In the ventricular cultures, *HAND1* and *FGF10* (associated predominantly with left- and right-ventricular development, respectively), *MYL6 (*smooth muscle), and *CDH2* (important for adherens junctions) were downregulated (Supplementary Table S2) (McFadden *et al*., 2005; Vega-Hernández *et al*., 2011). Very few genes were upregulated in ventricular cultures upon treatment with NMU. These findings further support that exogenous addition of NMU negatively impacts the efficiency of vCM differentiation. Notwithstanding a normal aCM differentiation efficiency in atrial cultures, downregulation of *WT1* and *BNC1* (both epicardial markers) and proliferation markers such as *MKI67* and *TOP2A* was observed, in addition to upregulation of *TTR*, a gene involved in transportation of vitamin A or retinol (Raghu and Sivakumar, 2004) (Figure 2F, Supplementary Table S1).

To further characterize the transcriptomic results, we performed GO-analysis on the up- and down-regulated gene sets found upon NMU treatment in the different cultures. This led to a limited number of enriched GO-terms (Figure S3) which generally are not heart-related. Next, we projected the combined expression levels of the gene lists back to the reference UMAP as presented in Figure 1B (Figure 2G). A compound score for each gene list was calculated for the wildtype cells, this allowed us to look at more global gene expression changes and relate those to the cell lineages present in the culture (Figure 1B). Notably, the atrial downregulated gene set upon NMU treatment, was specific to the cluster of epicardial cells. This corroborates the finding that both *WT1* and *BNC1* were downregulated upon NMU treatment and that this effect extends beyond those two examples of known epicardial markers. Upregulated genes after NMU treatment of the atrial cultures, showed expression in both aCM but also vCM progenitors branches of the UMAP, indicating NMU treatment affected genes that are not specific to either subtype, but broadly expressed across CMs. Under these conditions, also an increased expression is visible in ‘Other’ cells, which are related to anterior endoderm subtypes (Snabel *et al*., 2025). The ventricular downregulated gene set is expressed throughout the CMs and their progenitors, but highest in the cells committed to the vCM subtype, which was also reflected by the downregulation of *HAND1* and *FGF10* (Fig. 2F, Supplementary Table S2). Overall, these transcriptomic results show that in RA-treated atrial cultures, NMU reduces epicardial gene expression, whereas in the absence of RA, NMU treatment of the cultures specifically downregulates genes specific to the vCM signature which, however, does not disrupt the beating of these ventricular cardiac organoids, but rather changes their properties (see below).

### Administration of ET-1 results in increased aCM differentiation efficiencies

Using the same approach, we tested the influence of ET-1, an agonist of endothelin receptors, on the CM cultures, again measuring the reporter fluorescence levels and transcriptomic differences. *EDNRA* encodes an endothelin 1 (*EDN1, ET-1*) receptor, which showed increased expression in ventricular cultures at intermediate (day 8-10) and late (day 21) time points, whereas the atrial cultures only showed an increase in expression at late timepoints (Figure 1C, E). At earlier time points (day 5, early progenitors), *EDNRA* expression is detectable, albeit at much lower levels, in both culture conditions (Figure 1E). Interestingly, expression of its ligand, *EDN1*, was detected in only a few *TNNT2*-negative cells in both atrial and ventricular cultures (Figure S1B), whereas *EDNRA* is detected in a much larger population of cells, including both aCM and vCM progenitors (Figure 1E). *EDNRA* has been reported to be expressed in a subpopulation of both first heart field (FHF) and *ISL1*-positive second heart field (SHF) progenitors, including prospective atrial populations (Asai *et al*., 2010). To test the possible role of EDNRA during lineage specification *in vitro,* ET-1 was administered at different time points in both atrial and ventricular differentiation protocols (Figure 3A, Figure S2A). We found that ET-1 treatment from day 4 to 7 resulted in a significant decrease in the vCM differentiation efficiency (79% in control to 55% upon ET-1 treatment), whereas the aCM differentiation significantly increased (62% in control to 78% with ET-1), as determined by flow cytometry measurements of the *NKX2-5*-eGFP and *NR2F2*-mCherry reporter signals (Figure 2D and 3C, control and ET-1 treatment respectively). All other tested time point ranges of ET-1 administration did not have a significant effect on CM differentiation efficiency (Figure S2B-C).

**Figure 3:**
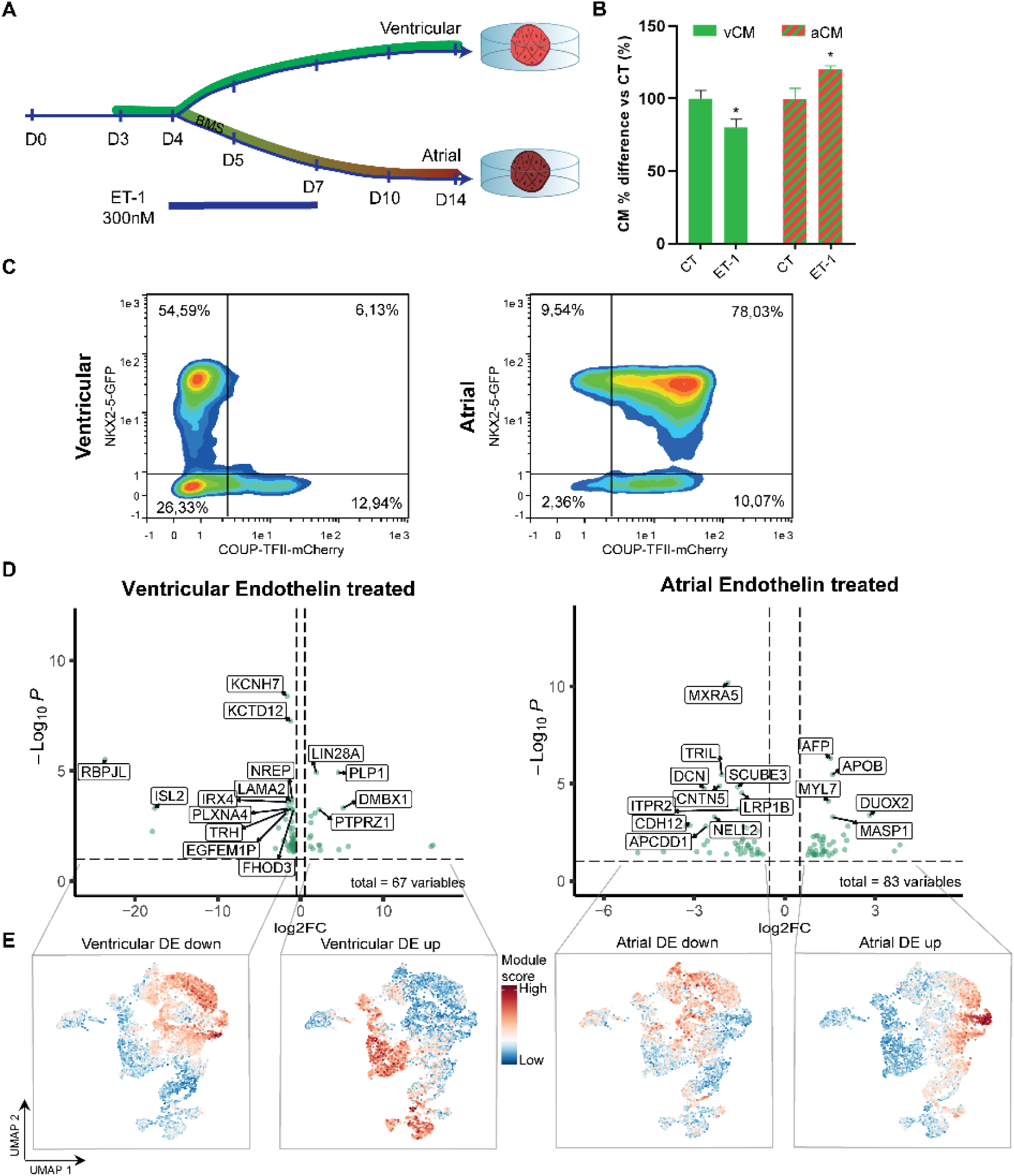
Transcriptomic analysis shows how endothelin treatment affects expression profiles associated with ventricular cardiomyocytes and uncovers lineage shifts due to signaling factor treatment. (**A**) Schematic representation of the culture conditions of the endothelin treated ventricular (upper) and atrial (bottom, RA-treated) protocols. (**B**) Average percentages of three replicates measuring the fluorescence of the NKX2-5-GFP and NR2F2-mCherry reporters as a proxy for ventricular (GPF-positive) and atrial (GPF- and mCherry-positive) cardiomyocytes in the culture after 14 days of culture and subjected to endothelin treatment (right bars) compared to control (CT, left bars). *p < 0.05. (**C**) Protein expression levels of NKX2-5-GFP on y-axis and NR2F2-mCherry on x-axis, as measured per event by means of FACS for cells under treatment with endothelin from day 4 until 7 of culturing. The control condition for this experiment is presented in Figure 2D. (**D**) Volcano plots with the significant (p < 0.05) differential expressed genes for the atrial (C) and ventricular (D) conditions treated with endothelin. (**E**) The EB reference UMAP (Figure 1B) labeled with compound gene expression levels (as module scores) based on the lists of genes retrieved from (for plots left to right): non-RA down- (log2FC < 0), non-RA up- (log2FC > 0), +RA down-, and +RA upregulated genes, based on significance (p-adjusted < 0.05) upon ET-1 treatment.

To gain further insights into the effects of administering ET-1 from day 4 to day 7 we analyzed the transcriptomes of the cells at day 14 along the two lineages (Figure 3D). The transcriptomic analysis at day 14 of the ventricular ET-1 treated culture, showed the significant downregulation of *ISL2* and the ventricular-associated genes *VCAM1, PLXNA4* and *IRX4* (Figure 3D, Supplementary Table S4), which was further confirmed with the gene ontology terms of cardiac-related processes enriched for this gene set (Figure S3), and with the population shifts observed with the FACS measurements (Figure S4A-B). These findings indicate that modulation of this pathway may adversely affect ventricular cardiac EBs. In atrial cultures, differential expression analysis between ET-1 treated and control samples, showed an upregulation of genes related to CMs, such as *HCN4*, *DES*, *PLN*, *MYLK3*, and the atrial-enriched *MYL7* gene, alongside an upregulation of the angiogenic factor *VEGFA* (Figure 3D, Supplementary Table S3). The epicardium marker *BNC1*, CM potassium channel *KCNJ2*, structural CM gene *COL6A3*, proliferation marker *MKI67*, protocadherin genes *PCDH11X* and *PCDH9,* and patterning factor *BMP4* were specifically downregulated in these atrial cultures. Gene ontology terms related to (striated) muscle cell development were found enriched in the differentially upregulated gene set (Figure S3), which is in line with the FACS observations that indicate a population shift towards more aCMs (Figure S4A-B). Overall, these results indicate an enhancing effect of ET-1 on the aCMs.

Broader patterns of gene expression, as observed with the compound expression levels of these gene lists (up- and down regulated genes upon ET-1 treatment of cultures treated with or without RA), further confirmed these patterns (Figure 3E). Treatment with ET-1 of the atrial cultures, resulted on day 14 of culture in the downregulation of genes which under control conditions are expressed in aCM progenitors and epicardial lineages. Interestingly, the upregulated genes in ET-1 treated atrial cells, are expressed specifically in the more late stage CMs (day 18-21). In the ventricular culture, an opposite pattern is observed, with an increase in early cardiac progenitor- and endodermal-enriched genes, and a downregulation of genes expressed in late CM progenitor branches. These results are in line with what was observed on reporter-based flow cytometry data, indicating a decrease of CMs in the ventricular cultures.

### Altered electrophysiology and signal propagation in both NMU and ET-1 treated cultures

To functionally test the effect of these lineage shifts occurring in the aCM and vCM cultures as a consequence of NMU or ET-1 treatment, we investigated electrophysiological properties of the cultures at day 14, with and without treatments (Figure 4). For the NMU treated cultures, we focused on the electrophysiology of ventricular cultures that have been treated for 24 hours with NMU, since that is where we observed the strongest effect. This revealed an increase in beating frequency (Figure 4A), to a rate that is similar to what normally is observed in atrial cultures. There were no significant differences in the action potential repolarization or conduction velocities (Figure 4B-C). These findings indicate that exogenous addition of NMU resulted in electrophysiological changes with a significantly increased beating frequency.

**Figure 4:**
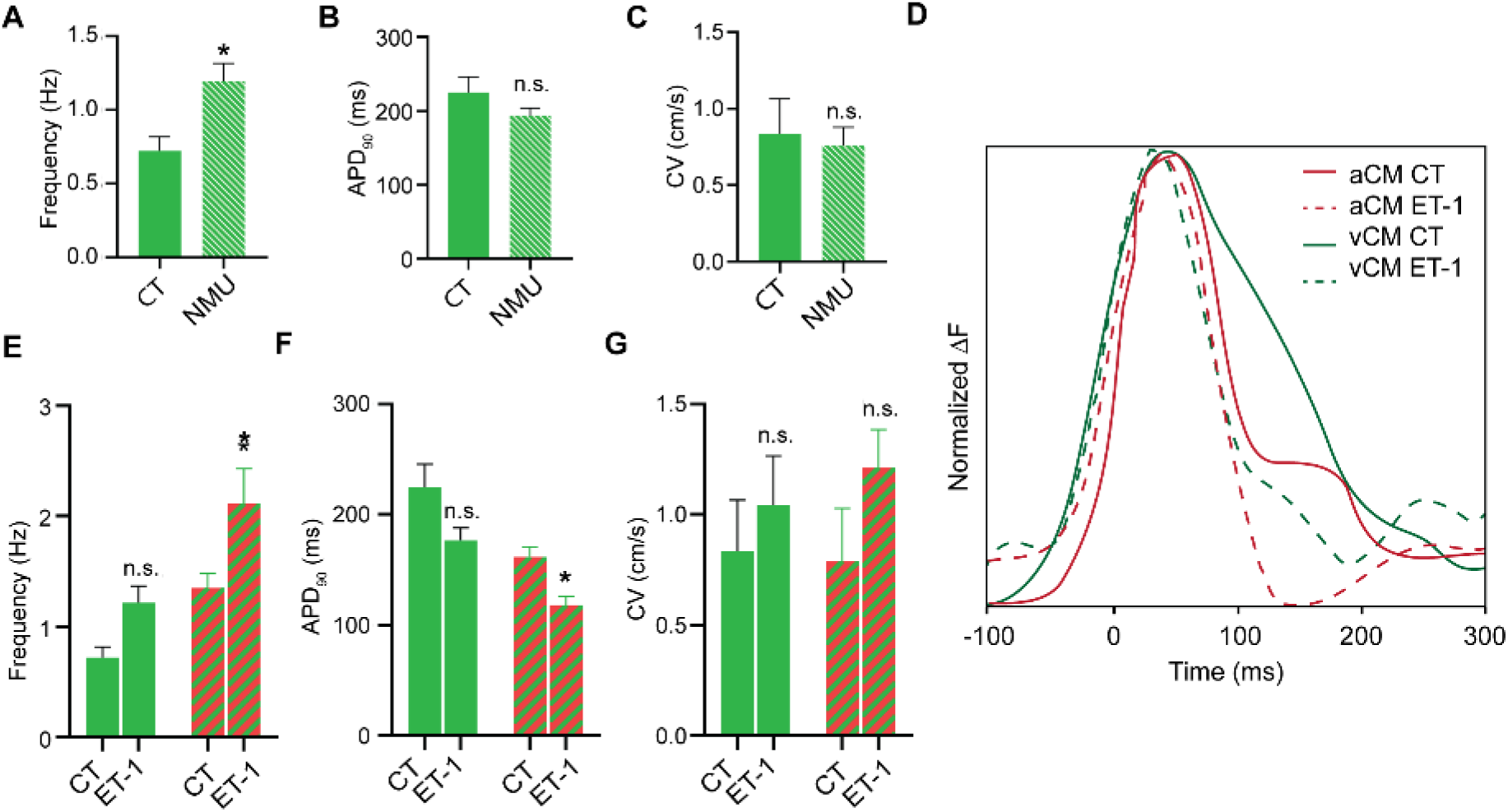
Physiological readout of treated cultures indicates that NMU increases beating frequency of ventricular cultures and ET-1 changes electrophysiological properties within the culture. (**A-C**) Effect of NMU treatment from day 4 to day 5 during ventricular EB differentiation evaluated by (A) the beating frequency (B) action potential duration at 90% of repolarization and conduction velocity (C) at day 14 compared to control (CT). (**D-F**) Effect of ET-1 treatment on the ventricular and atrial cultures, with in (D) the representative action potential duration, in (E) the frequency of beating, in (F) Quantification of APD90 upon ET-1 treatment. (**G**) Conduction velocity in atrial or ventricular cultures treated with ET-1 or without (CT).

Action potential maps indicated a different electrical activation pattern of the contraction between controls and ET-1 treated aCMs, where controls showed a starting point of the action potential in one edge of the EB from where it propagated towards the other end, and the ET-1-treated aCMs exhibiting a rotating-like pattern of the action potential, travelling around the edges of the EB (Figure S4 and Videos S1-6). Contraction frequency was significantly increased in the aCM cultures upon treatment with ET-1 (Figure 4D-E). Moreover, ET-1-treated aCMs demonstrated significantly shorter repolarization times (APD90) compared to the control group (Figure 4F). Despite the rotating patterns of action potential progression, ET-1 treatment did not correlate with differences in conduction velocity (Figure 4G). These results suggest that ET-1 treatment not only causes differences in the number of atrial and ventricular CMs, but is also associated with an increase in contraction frequency that may be caused by more rapid repolarization and a shorter refractory period. These findings together indicate an increase in the number of atrial CMs, with an additional change in action potential progression inside these cardiac organoids cultures, suggesting that ET-1 treatment of atrial cultures from the cardiac mesoderm stage until the CM progenitor stage contributes to an overall improvement of atrial differentiation with altered electrophysiological properties.

## Discussion

To focus on the signaling differences between the atrial and ventricular lineage across the 21-day cardiac EB protocol, we analyzed the transcriptomes of cells differentiated with or without RA administration (atrial and ventricular cultures). Through the direct comparison of these EB-derived CM subtypes, we identified signaling factors predicted to have the potential to modulate cardiac lineage commitment. We showed how the single-cell time series data can be utilized to specifically probe the roles of signaling factors, and the lineage commitment shifts that they can induce. We identified *EDNRA* and *NMU* among the signaling factors and receptors with an unknown role in cardiac lineage specification, and after administering specific agonists for these signaling pathways, we performed a detailed exploration of the influence of these signals on lineage progression and CM differentiation efficiency. The results uncover unanticipated positive and negative quantitative effects of these signals on subtype-specific cardiomyocyte traits, highlighting the importance of signal-modulation in cardiac organoid cultures.

NMU is known to exhibit vasoconstrictive properties and acts on smooth muscle cells (Minamino *et al*., 1985; Maggi *et al*., 1990), and it is also more abundant in blood samples of heart failure patients (Healy *et al*., 2024). Not much is known about the role of NMU in heart development. Our results uncover a potential role for the NMU pathway in early cardiac lineage commitment. In NMU-treated atrial cultures, bulk RNA-seq showed decreased expression of epicardial- and ventricular-associated genes, suggesting decreased numbers of epicardial and vCMs. Although aCM differentiation efficiency was not substantially increased, our findings indicate that NMU could have an atrial CM- enhancing function. The specific expression of NMU around day 7-8 of culturing coincides with the time at which cardiac cell lineages begin to diverge in these cultures, as previously shown in the single-cell roadmap (Snabel *et al*., 2025). In ventricular cultures, we found that NMU increased beating frequency. This may be related to gene expression changes, including a decreased *CDH2* expression. Reduced CDH2 expression has been linked to electrical uncoupling and increased arrhythmia susceptibility *in vivo*, through disruption of intercalated disc organization and connexin-based gap junctions (Li *et al*., 2005). Moreover, mutations in *CDH2* have been found to cause ventricular arrhythmogenic cardiomyopathy (Mayosi *et al*., 2017; Turkowski *et al*., 2017; Ghidoni *et al*., 2021). Although our cultures did not show a significant decrease in conduction velocity, the increased beating frequencies observed following NMU treatment of our ventricular cultures, could possibly reflect an intrinsically increased beating rate or a less synchronized electrical activity. Further experiments will be required to distinguish between these possibilities. Together with the downregulation of *NEXN* and *SMPX*, which are involved in the cytoskeletal organization, and *MYL6*, which is associated with actomyosin machinery, these changes suggest a remodeling of the structural and contractile program in these cultures. In CMs, intercalated disc and cytoskeletal organization are closely linked to electrical coupling and synchrony (Nielsen *et al*., 2023). Furthermore, downregulation of *HAND1* upon NMU treatment, an important cardiac transcription factor for ventricular specification, may indicate a shift away from a ventricular-like state. Since atrial and ventricular CMs differ in their electrophysiological and structural properties, these results indicate that NMU modifies the differentiation trajectory of cardiac progenitors. However, further investigation is required to determine whether NMU directly promotes atrial specification or instead alters ventricular maturation and electrical coordination.

Endothelin (ET-1) encoded by the *EDN1* gene is predominantly expressed in the endothelial cells of fetal and adult hearts (Knight-Schrijver *et al*., 2026), but it is not expressed in the endothelial cells in our cardiac cultures (Suppl. Fig. S1). The differential response between lineages to treatment with ET-1, may be linked to *EDNRA* expression patterns. *EDNRA* is expressed in the cardiac crescent and plays a key role in myocardial chamber formation (Asai *et al*., 2010). In our system, EDNRA was present in both lineages from day 5, with significant upregulation in ventricular cultures from day 8. Conversely, atrial cultures did not show increased *EDNRA* expression until day 21 of differentiation. Expression of the receptor in the atrial progenitor populations during heart tube stages of development has been reported (Sedmera, 2014). We hypothesized that early administration of ET-1 could enhance the development of CMs because of the presence of this receptor, while *EDN1* expression was not detected at early time points. Triggering this endothelin pathway had most impact in the atrial cultures, leading to an increased aCM differentiation efficiency. This was accompanied by an upregulation of late CM-specific genes and altered electrophysiological parameters, specifically increased frequency and faster repolarization. These findings suggest that addition of ET-1 during early stages of differentiation promotes a more mature CM cellular state and that modulation of the ET-1/*EDNRA* pathway may play a significant role in atrial specification. These effects on differentiation were markedly CM subtype-specific, as ET-1 modulation in ventricular cultures reduced the number of vCMs, suggesting important crosstalk of RA and ET-1. This warrants further investigation. Moreover, several studies have reported that exogenous ET-1 can induce a switch in the differentiation towards myoconductive cell types during chick and mouse embryo development, both in *in vivo* and *in vitro* settings (Gourdie *et al*., 1998; Patel and Kos, 2005; Sedmera, 2014). Although we did not observe this transition in our culture based on the specific markers of these cells, the atrial cultures did show signs of increased myoconduction by, for example, a faster repolarization.

In summary, our results provide a proof of principle approach in studying the signal transduction pathways involved in atrial and ventricular cardiac lineages, using a combination of single-cell data, perturbation experiments and bulk transcriptomic follow-up analysis. The data from the perturbation experiments can be mapped back onto the single-cell roadmap reference, in exploration of lineage shifts as a consequence of these perturbations. Our results indicate how the plasticity of these cardiac cultures was exploited by the administration of external signals targeting the NMU and Endothelin pathways. The presented findings provided indications for a role of NMU in increasing the beating frequency in ventricular cardiac embryoids, as well as a role for ET-1 in increasing atrial CM differentiation efficiency and reducing the repolarization time in RA-induced cultures. Further characterization of these signaling pathways in these cultures will be needed to underpin the molecular mechanisms affected downstream of the pathways, however, our work showcases how our set-up can play a role in further exploring these molecular principles.

## Methods

### Single-cell temporal roadmap construction and EB selection

The cells harvested from the EB cultures were selected from the roadmap as previously described (Snabel *et al*., 2025), this paper details the steps of hPSC-derived cardiomyocyte EB cultures, scRNA-seq library preparation and the mapping of the data. In short, the cardiac EB cultures were generated using the human embryonic stem cell (hESC) lines NKX2.5^EGFP/+^-COUP-TFII^mCherry/+^ (Schwach *et al*., 2017) and differentiated into ventricular or atrial EBs, using the protocol as previously described (Schwach *et al*., 2022) for a total of 21 days. On day 4, atrial EBs were refreshed with 20 µM RARα selective agonist BMS-753 (Torcis Bioscience, 3505). The plate-based scRNA-seq libraries were generated with a modified SORT-seq protocol according to protocol (Hashimshony *et al*., 2016; Muraro *et al*., 2016). The primers used to enrich poly-A mRNAs were designed reversed from classic CELSeq2 primers (as described in Gerlach *et al*., 2019 (Gerlach *et al*., 2019)). Sequencing was performed with the NextSeq 500 (Illumina). Mapping of the reads was performed with kb-python v0.25.0 (kb count with the setting ‘--technology 1,8,16:1,0,8:0,0,0’) against the GRCh38 genome, which quantified the UMI counts per gene per cell (Melsted *et al*., 2021). From the total roadmap produced with these steps, only the cells harvested from EB culture experiments were kept and combined into one dataset, as described for the transcription factor selection in the EB subset of the roadmap. The cells derived from separate EB differentiation experiments were integrated using Seurat v3, as described before (Snabel *et al*., 2025).

### Optical membrane potential imaging

For optical membrane potential imaging, aCMs or vCMs at day 14 were stained for 30 min at RT with the voltage-sensitive dye FluoVolt™ according to manufacturer. For signal acquisition, the loaded sample was excited using a 488nm laser and recorded for 10 seconds. Movies were acquired with a Nikon ECLIPSE Ti2 fluorescent microscope under temperature and humidity control (37°C and 5% CO_2_). Videos were analyzed with BV Workbench, Brainvision Inc.

### FACS and flow cytometry

Day 3 to day 8 EBs were dissociated to single cells with TypLE 1X (Thermo Fisher, 12563029) for 5 minutes at 37°C. From day 8 onwards, the EBs were dissociated with TypLE 10X (Thermo Fisher, A1217702) for 10 min at 37°C. Cells were sorted as single cells into 384-well primer plates for single-cell RNA-sequencing. Sorting was performed with a Sony SH800. Flow cytometry was performed with a MACSQuant Analyzer 16 (Miltenyi). Flow cytometry data was analyzed with FlowJo software (Miltenyi).

### Modulation of EB protocol with signaling factors of interest

Endothelin 1 (ET-1) treatment (0.3 µM, Sigma Aldrich, E7764) was performed through the atrial and ventricular EB differentiation protocols. Several timepoints were tested: administration from day 4 to 5, from day 4 to 7, from day 4 to 14, from day 5 to 7, or from day 10 to 14 (Supplementary Figure S2A). If ET-1 treatment fell in a refreshment timepoint of the differentiation protocol, such as day 6, new ET-1 was administered. Neuromedin U (NMU) modulation was tested in atrial and ventricular EB differentiation protocols by administration of 10 nM NMU (Abcam, ab141007) at different timepoints: from day 4 to 5, from day 4 to 6, or from day 4 to 7 (Supplementary Figure S2A).

### Bulk RNA-sequencing sample preparation

RNA was harvested at day 14 of the protocols and purified using the Nucleo Spin RNA (Macherey-Nagel) kit according to the manufacturer’s protocol. Libraries were generated with the RiboErase depletion RNA-sequencing protocol by KAPA (Roche, 8098131702) and sequenced on the Illumina Next Seq 2000. Per sample 200 ng of total RNA was used for library preparations with the KAPA RNA HyperPrep Kit with RiboErase (HMR) (Roche) as instructed by the manufacturer. A total of three replicates per differentiation condition were taken along, except for RA- and NMU-treated, which consisted of two replicates.

## Computational analysis

### Differential signaling factor analysis and module scores

For direct aCM and vCM comparison, a subset of only embryoid body harvest time points were selected from the larger temporal atlas (detailed in (Snabel *et al*., 2025)). Differential gene expression testing was performed on selected clusters, representing the progenitor stages contributing to aCM and vCM over time. Wilcoxon rank-sum test was used to find significant differentially expressed genes between the clusters 24 and 6, 14 and 0, 9 and 4 and 2 and 3 (atrial and ventricular clusters, respectively, according to the clustering described before, and shown in Figure 3 of the paper (Snabel *et al*., 2025)), expressed in at least 25% of the cells in one of the clusters in each comparison and with an adjusted p-value < 0.05. The resulting gene lists were filtered for genes present in the CellPhoneDB v2 database (Efremova *et al*., 2020), to solely obtain signaling factors among the differentially called genes. Hierarchical clustering of the signaling factors was performed to separate specific expression patterns over the different time points across the lineages, and the results were visualized using Complex Heatmap v2.8.0 (Gu, Eils and Schlesner, 2016). Module scores across the EB atlas were calculated using the ModuleScore() functionality in Seurat, with the sets of differential genes up- or downregulated in the bulk transcriptomic data obtained at day 14 of ventricular and atrial culture, after NMU or ET-1 treatment (Table S1-4).

### Validation screens RNA-seq mapping and differential analysis

The Seq2Science v0.9.6 pipeline was used to map the RNA-seq reads to the GRCh38 genome (van der Sande *et al*., 2023). Differential gene expression analysis between control and targeted conditions was performed with the R package DESeq2 (v1.34.0) (Love, Huber and Anders, 2014). The differential gene list was filtered for a log2 fold change > 1 and P-adjusted value < 0.05, and visualized with pheatmap, EnhancedVolcano v1.12.0 (Blighe, Rana and Lewis, 2019) and clusterProfiler v4.2.0 (Wu *et al*., 2021). The counts per million (CPM) were log1p-transformed.

## Author contributions

R.R.S. conceived experiments, generated libraries for RNA-sequencing experiments, led the computational analysis and wrote the manuscript. C.C.F. conceived experiments, performed all cell culture work, harvested and sorted the cells, performed perturbation experiments and wrote the manuscript. M.B. generated bulk RNA-sequencing libraries and performed the sequencing. V.S. conceived experiments. R.P. conceived experiments, supervised the experiments involving stem cell differentiation, revised the manuscript and generated funding for the work. G.J.C.V. conceived experiments, supervised computational genomics analysis, revised the manuscript and generated funding for the work.

## Acknowledgements

This work was funded by ZonMw TOP (project number 91217061) and NWO XL (dossier OCENW.XL21.XL21.067, HeartEngine) grants to R.P. and G.J.C.V. The authors thank all HeartEngine consortium members for helpful discussions.

## Declaration of interest

R.P. is a cofounder of Pluriomics (Ncardia) and River BioMedics BV. The other authors declare no competing interests.

---

**Supplementary Figure S1:**
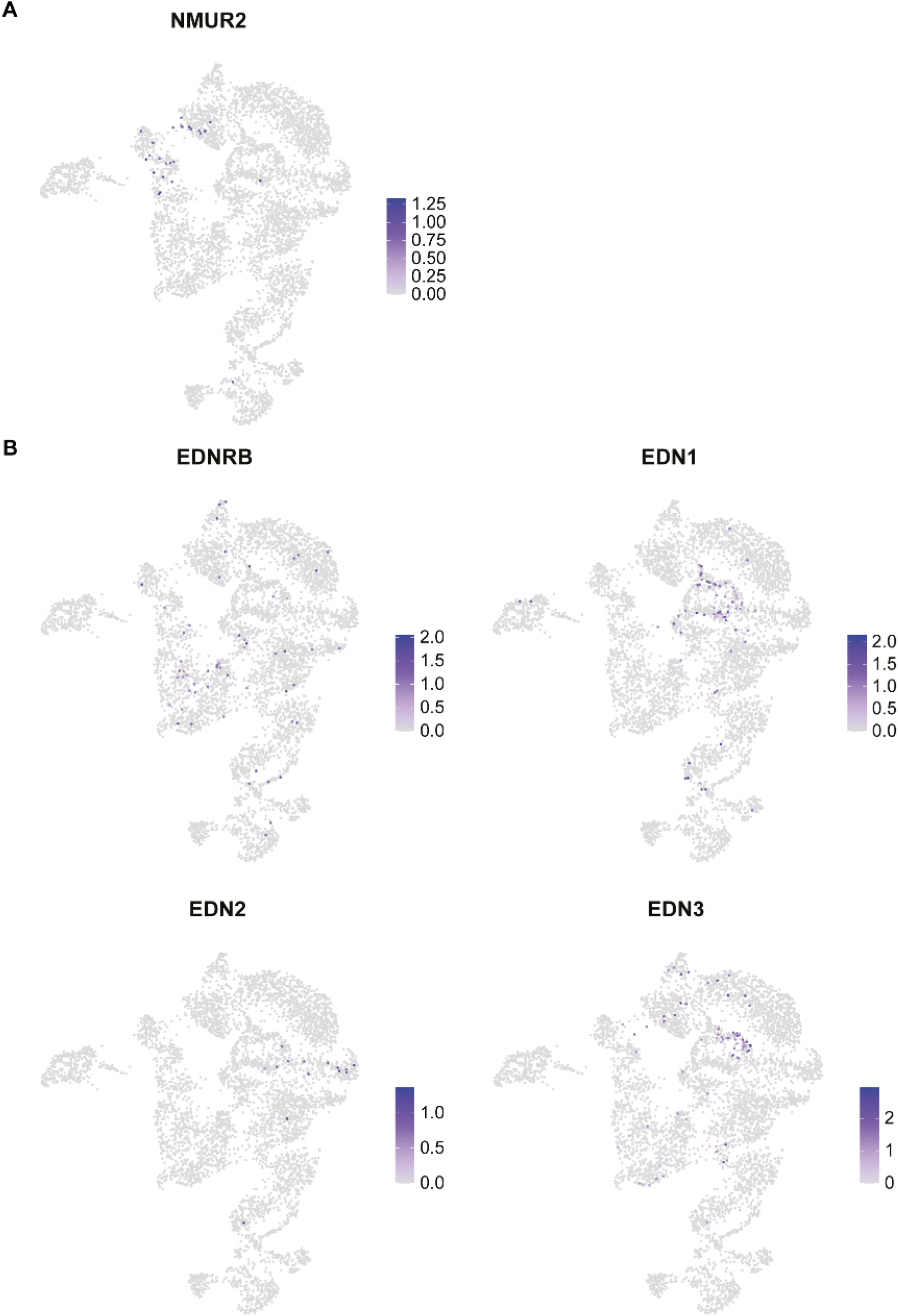
Expression across the scRNA-seq dataset of receptors and ligands related to the two selected signaling factors of interest, EDNRA and NMU. (A) Expression levels of another endothelin receptor, EDNRB and three of the ligands EDN1, EDN2 and EDN3. (B) Expression levels of the NMU receptor 2.

**Supplementary Figure S2:**
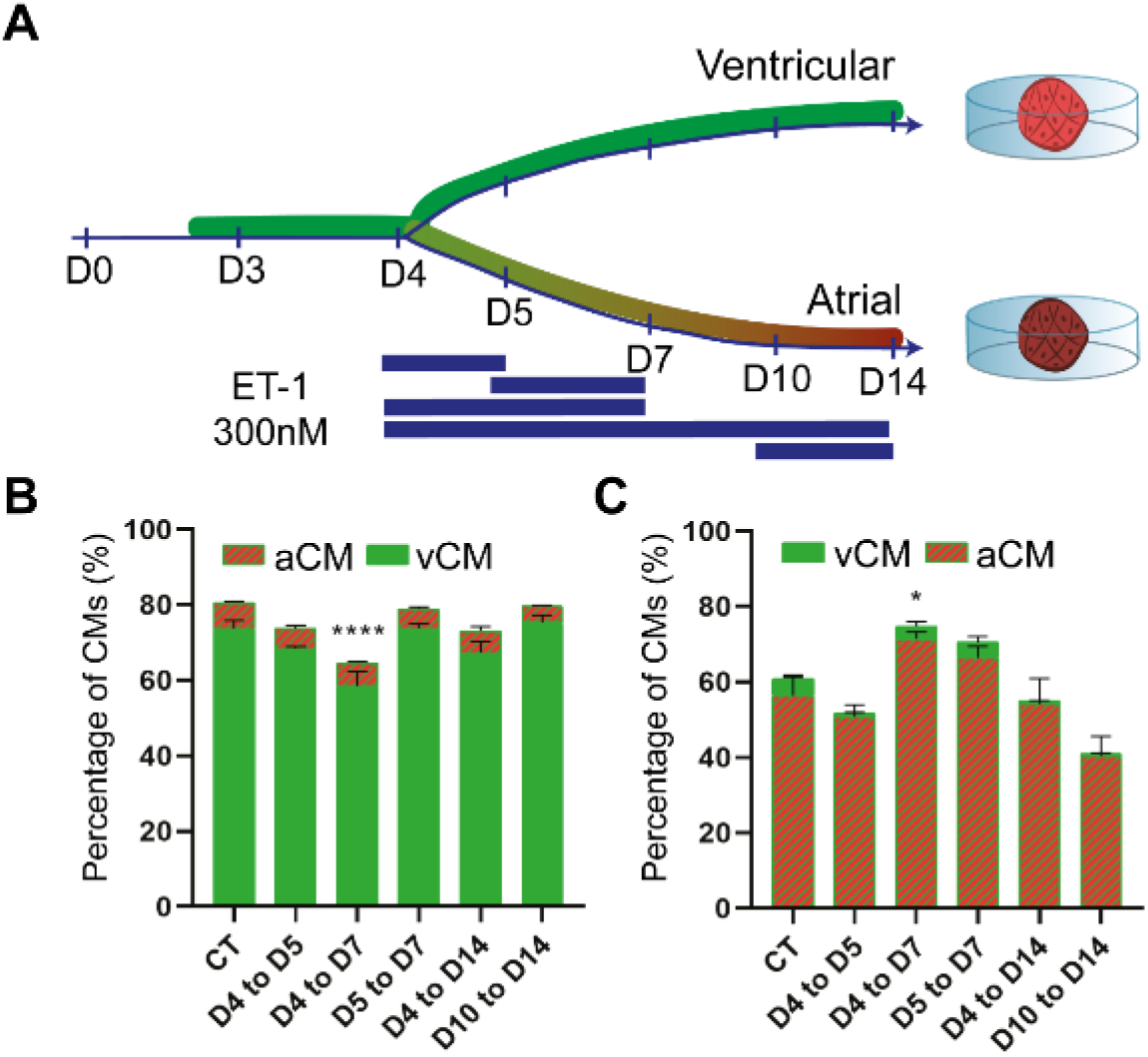
Exploration of different ET-1 treatment timings during ventricular and atrial cardiomyocyte differentiation protocols. (A) Schematic overview of the culture with indication of different timing of external ET-1 explored. (B-C) Percentage of reporters (aCM = NKX2-5- and NR2F2-positive, vCM = NKX2-5-positive and NR2F2-negative) as measured with FACS for the different ET-1 treatment timings versus no treatment (CT) in ventricular culture (B) and atrial culture (C), measured in triplicate.

**Supplementary Figure S3.**
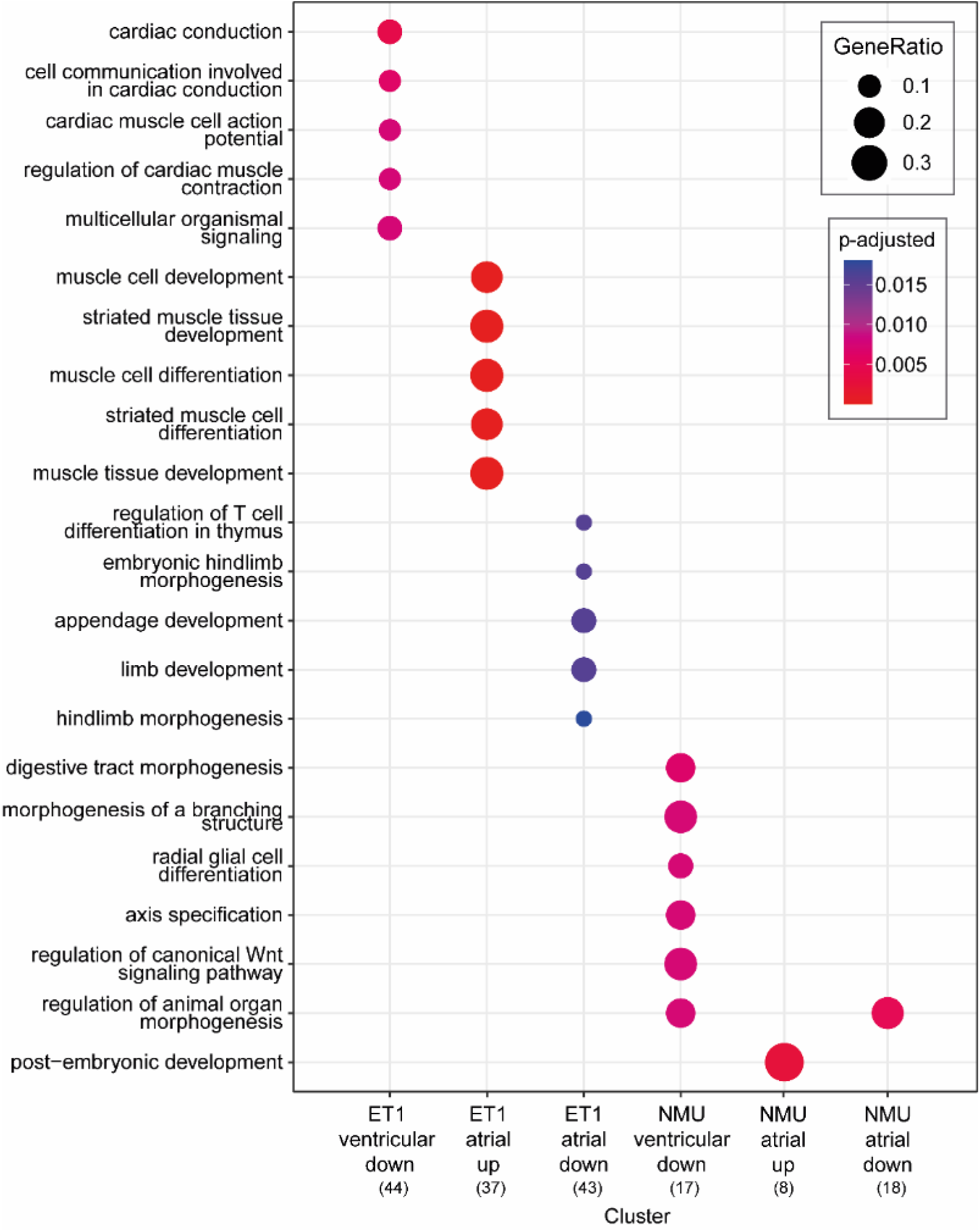
: Gene ontology analysis of differentially upregulated or downregulated genes, after ET-1 or NMU treatment of a ventricular and atrial cardiomyocyte culture. In parenthesis the number of genes within the cluster used for gene ontology (GO) enrichment analysis, are indicated.

**Supplementary Figure S4:**
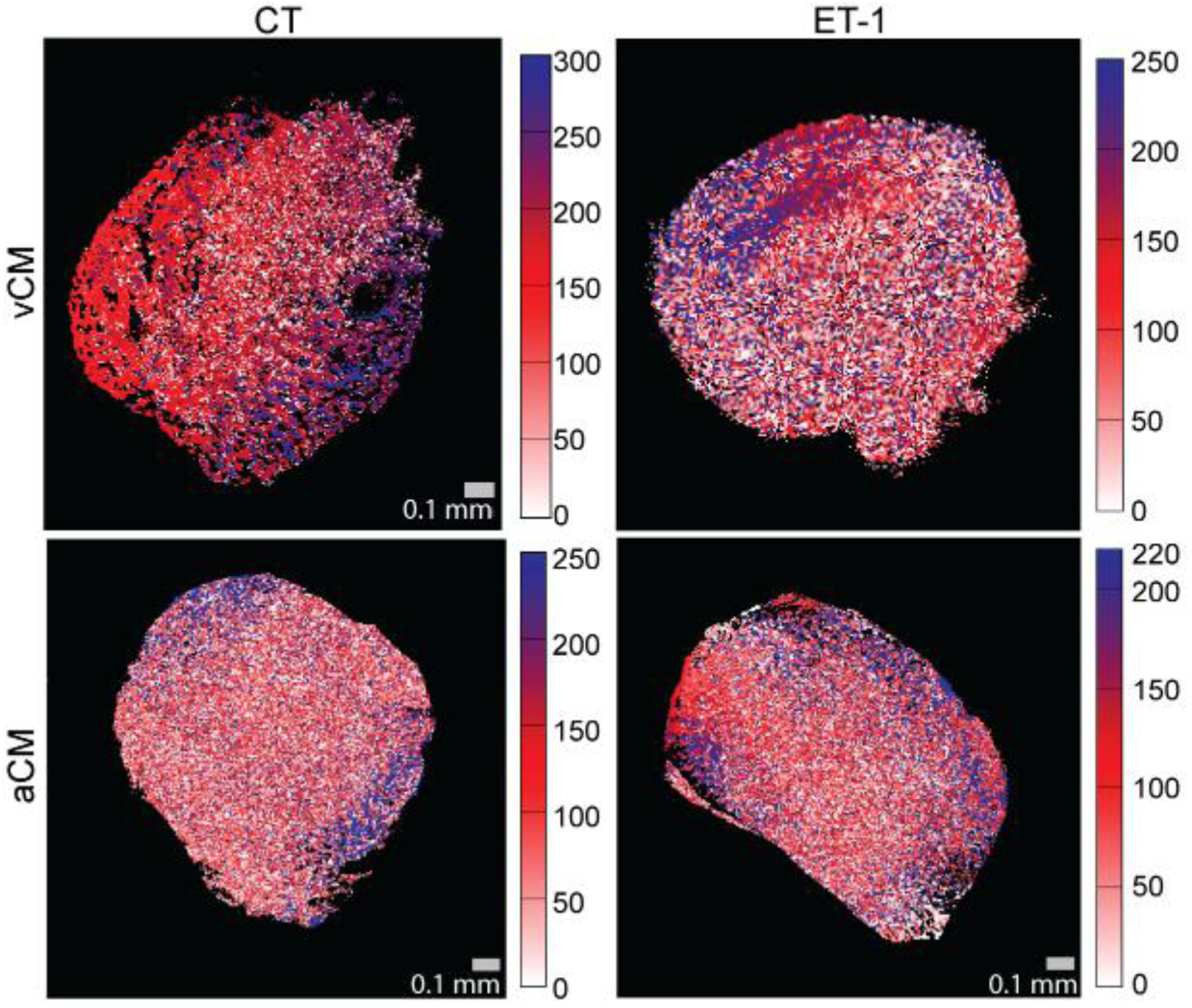
Action potential propagation maps after ET-1 treatment of ventricular and atrial CM cultures. The representative activation maps from FluoVolt measurement in control (CT) and ET-1 atrial and ventricular EBs at day 14. Scale bar = 1 mm. Gradient bar represents time in ms.

**Supplementary Table S1.** : Atrial NMU treatment over control, differential genes

|  | baseMean | log2FoldChange | lfcSE | stat | p-value | p-adjusted |
| --- | --- | --- | --- | --- | --- | --- |
| WT1 | 26.66325 | -5.22355 | 1.326174 | -3.93881 | 8.19E-05 | 0.036735 |
| HGF | 38.38822 | -5.0028 | 1.037741 | -4.82085 | 1.43E-06 | 0.002084 |
| FGF5 | 24.98867 | -4.08298 | 1.012957 | -4.03075 | 5.56E-05 | 0.027021 |
| SBSPON | 52.22484 | -3.27118 | 0.673592 | -4.85632 | 1.20E-06 | 0.001993 |
| DCN | 57.57465 | -2.7949 | 0.546702 | -5.11229 | 3.18E-07 | 0.000947 |
| BNC1 | 134.4849 | -1.88583 | 0.382936 | -4.92465 | 8.45E-07 | 0.001643 |
| KIF11 | 87.92572 | -1.79453 | 0.434067 | -4.13422 | 3.56E-05 | 0.021865 |
| CXCL12 | 305.6214 | -1.75759 | 0.435475 | -4.03602 | 5.44E-05 | 0.027021 |
| ANKRD1 | 366.0845 | -1.64826 | 0.325342 | -5.06623 | 4.06E-07 | 0.000947 |
| CCN1 | 377.6305 | -1.6108 | 0.28965 | -5.56118 | 2.68E-08 | 0.000156 |
| LGI2 | 181.6209 | -1.57976 | 0.346835 | -4.55477 | 5.24E-06 | 0.004705 |
| CDH11 | 350.1621 | -1.51581 | 0.321364 | -4.7168 | 2.40E-06 | 0.002809 |
| CCN2 | 397.0324 | -1.49681 | 0.339122 | -4.41379 | 1.02E-05 | 0.008463 |
| ENC1 | 469.7314 | -1.45376 | 0.308755 | -4.70845 | 2.50E-06 | 0.002809 |
| DAB2 | 156.0343 | -1.35806 | 0.336913 | -4.0309 | 5.56E-05 | 0.027021 |
| MKI67 | 474.8936 | -1.32815 | 0.288562 | -4.60265 | 4.17E-06 | 0.004055 |
| TOP2A | 283.886 | -1.31087 | 0.318927 | -4.11025 | 3.95E-05 | 0.022886 |
| GPC6 | 312.6708 | -1.2178 | 0.296979 | -4.10062 | 4.12E-05 | 0.022886 |
| RAI14 | 224.9443 | -1.21455 | 0.309299 | -3.92679 | 8.61E-05 | 0.03719 |
| VEGFA | 1417.398 | 1.085883 | 0.253817 | 4.278216 | 1.88E-05 | 0.013734 |
| AFP | 1166.336 | 1.090481 | 0.27365 | 3.984947 | 6.75E-05 | 0.031491 |
| CKB | 809.2357 | 1.491131 | 0.317509 | 4.696337 | 2.65E-06 | 0.002809 |
| APOB | 672.4562 | 1.545496 | 0.304488 | 5.075722 | 3.86E-07 | 0.000947 |
| TRIB3 | 133.7768 | 1.580676 | 0.381551 | 4.142763 | 3.43E-05 | 0.021865 |
| UPP1 | 281.8442 | 1.922662 | 0.458129 | 4.196773 | 2.71E-05 | 0.018576 |
| PPP1R13L | 196.165 | 1.92957 | 0.446686 | 4.31975 | 1.56E-05 | 0.012147 |
| TTR | 459.3009 | 2.24658 | 0.396227 | 5.669932 | 1.43E-08 | 0.000156 |

**Supplementary Table S2.** : Ventricular NMU treatment over control, differential genes

|  | baseMean | log2FoldChange | lfcSE | stat | p-value | p-adjusted |
| --- | --- | --- | --- | --- | --- | --- |
| PERCC1 | 1.022123 | -23.706 | 3.583011 | -6.61622 | 3.69E-11 | 5.75E-07 |
| LGSN | 0.728078 | -23.0929 | 3.999685 | -5.77368 | 7.76E-09 | 4.03E-05 |
| ADH1A | 0.530105 | -17.4695 | 4.025442 | -4.33978 | 1.43E-05 | 0.015215 |
| C22orf23 | 0.634758 | -17.2152 | 3.997446 | -4.30656 | 1.66E-05 | 0.015215 |
| TDGF1 | 42.56555 | -2.39613 | 0.534911 | -4.47949 | 7.48E-06 | 0.01061 |
| SCARNA7 | 118.1038 | -1.4079 | 0.319482 | -4.40683 | 1.05E-05 | 0.012587 |
| RSPO3 | 558.8669 | -1.33413 | 0.256445 | -5.20239 | 1.97E-07 | 0.000438 |
| MT-ND3 | 755.7655 | -1.30906 | 0.237098 | -5.52116 | 3.37E-08 | 9.15E-05 |
| HAND1 | 273.542 | -1.2998 | 0.275128 | -4.72434 | 2.31E-06 | 0.004001 |
| RMRP | 4296.312 | -1.25856 | 0.277341 | -4.53797 | 5.68E-06 | 0.00886 |
| MT-ND4L | 366.5301 | -1.14458 | 0.284288 | -4.02613 | 5.67E-05 | 0.038456 |
| KCTD12 | 2567.873 | -1.0811 | 0.178964 | -6.04088 | 1.53E-09 | 1.20E-05 |
| SMPX | 323.3594 | -0.98498 | 0.247998 | -3.97174 | 7.13E-05 | 0.042806 |
| NREP | 2801.726 | -0.98171 | 0.193094 | -5.08413 | 3.69E-07 | 0.00072 |
| PKIA | 374.835 | -0.91793 | 0.207709 | -4.41929 | 9.90E-06 | 0.012587 |
| NEXN | 482.9835 | -0.9134 | 0.232196 | -3.93376 | 8.36E-05 | 0.048315 |
| FGF10 | 361.7036 | -0.87207 | 0.207084 | -4.21118 | 2.54E-05 | 0.020856 |
| SFRP1 | 1119.771 | -0.86954 | 0.201715 | -4.31075 | 1.63E-05 | 0.015215 |
| MYL6 | 1128.292 | -0.82472 | 0.193217 | -4.26837 | 1.97E-05 | 0.017064 |
| CDH2 | 2609.407 | -0.70201 | 0.173178 | -4.05371 | 5.04E-05 | 0.037447 |
| SCNN1A | 56.03398 | 1.707707 | 0.417341 | 4.091873 | 4.28E-05 | 0.033374 |
| XACT | 720.2263 | 1.738839 | 0.307554 | 5.653762 | 1.57E-08 | 6.12E-05 |
| MIAT | 91.67659 | 1.891346 | 0.46894 | 4.033236 | 5.50E-05 | 0.038456 |
| OTX1 | 62.14143 | 2.505368 | 0.580664 | 4.31466 | 1.60E-05 | 0.015215 |
| TLL1 | 26.72728 | 2.602806 | 0.651877 | 3.992784 | 6.53E-05 | 0.040746 |
| MAP3K7CL | 8.355448 | 5.917451 | 1.479325 | 4.000102 | 6.33E-05 | 0.040746 |
| MAD2L1P1 | 3.078773 | 21.55959 | 3.91042 | 5.513369 | 3.52E-08 | 9.15E-05 |

**Supplementary Table S3.**
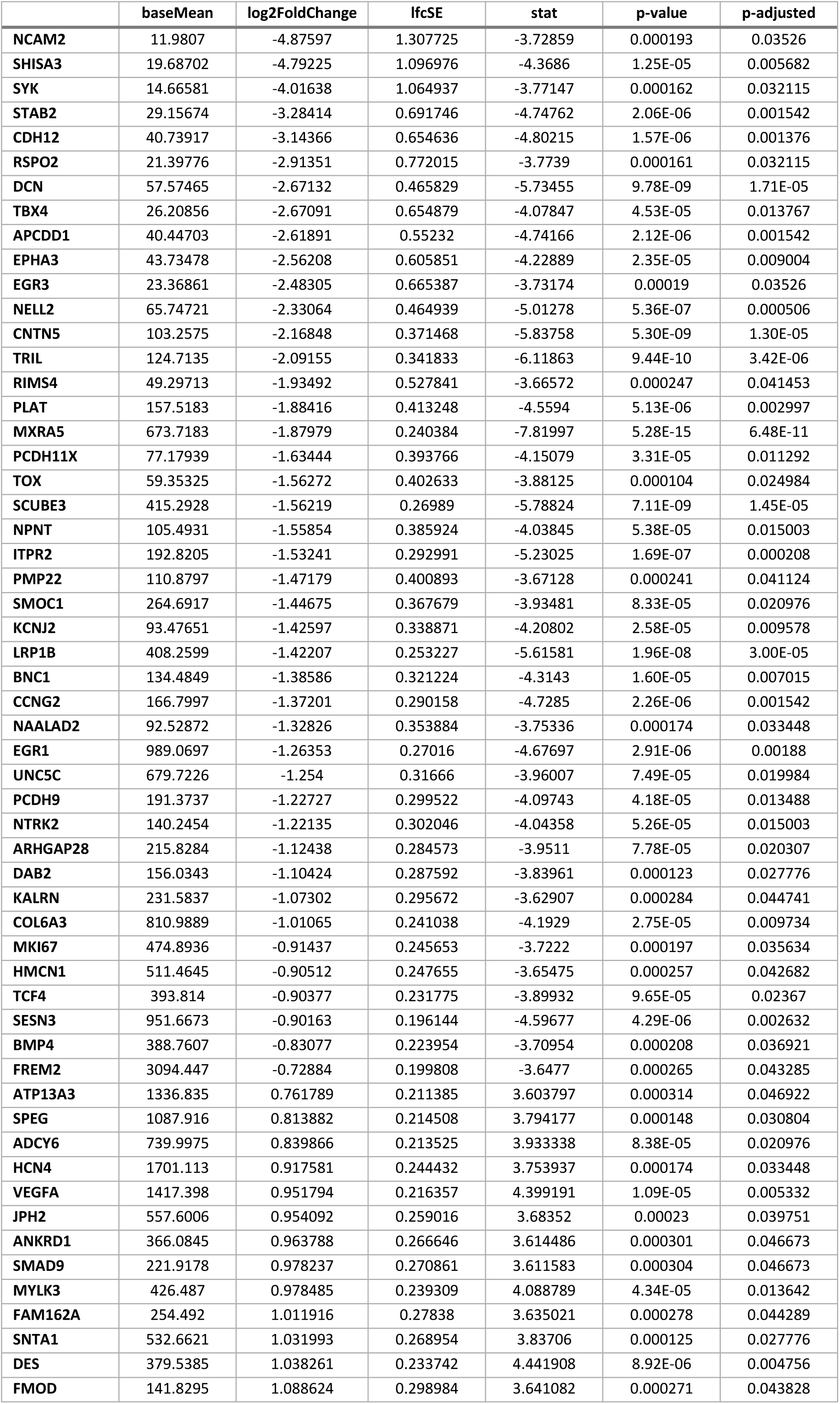
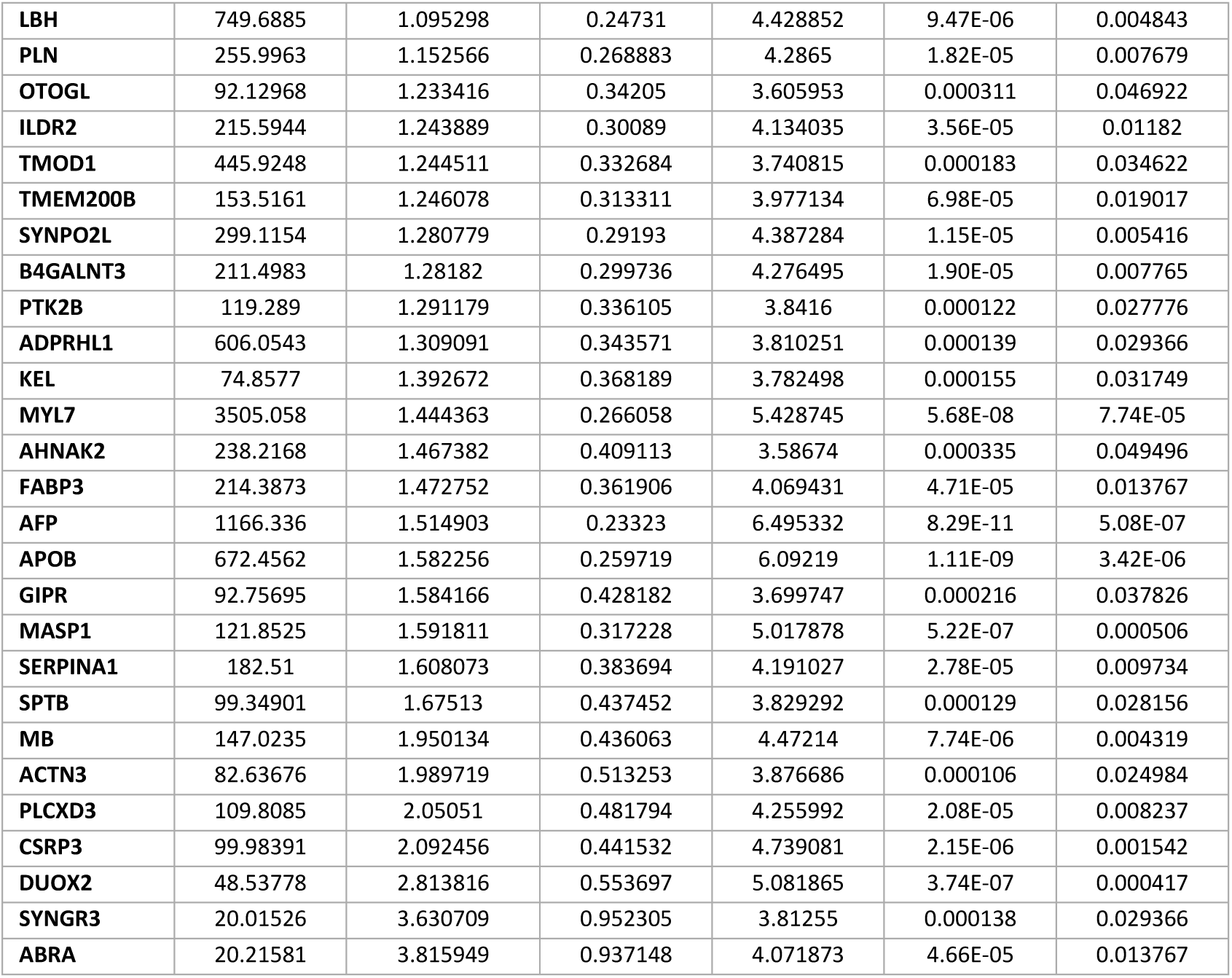
: Atrial ET-1 treatment over control, differential genes

|  | baseMean | log2FoldChange | lfcSE | stat | p-value | p-adjusted |
| --- | --- | --- | --- | --- | --- | --- |
| NCAM2 | 11.9807 | -4.87597 | 1.307725 | -3.72859 | 0.000193 | 0.03526 |
| SHISA3 | 19.68702 | -4.79225 | 1.096976 | -4.3686 | 1.25E-05 | 0.005682 |
| SYK | 14.66581 | -4.01638 | 1.064937 | -3.77147 | 0.000162 | 0.032115 |
| STAB2 | 29.15674 | -3.28414 | 0.691746 | -4.74762 | 2.06E-06 | 0.001542 |
| CDH12 | 40.73917 | -3.14366 | 0.654636 | -4.80215 | 1.57E-06 | 0.001376 |
| RSPO2 | 21.39776 | -2.91351 | 0.772015 | -3.7739 | 0.000161 | 0.032115 |
| DCN | 57.57465 | -2.67132 | 0.465829 | -5.73455 | 9.78E-09 | 1.71E-05 |
| TBX4 | 26.20856 | -2.67091 | 0.654879 | -4.07847 | 4.53E-05 | 0.013767 |
| APCDD1 | 40.44703 | -2.61891 | 0.55232 | -4.74166 | 2.12E-06 | 0.001542 |
| EPHA3 | 43.73478 | -2.56208 | 0.605851 | -4.22889 | 2.35E-05 | 0.009004 |
| EGR3 | 23.36861 | -2.48305 | 0.665387 | -3.73174 | 0.00019 | 0.03526 |
| NELL2 | 65.74721 | -2.33064 | 0.464939 | -5.01278 | 5.36E-07 | 0.000506 |
| CNTN5 | 103.2575 | -2.16848 | 0.371468 | -5.83758 | 5.30E-09 | 1.30E-05 |
| TRIL | 124.7135 | -2.09155 | 0.341833 | -6.11863 | 9.44E-10 | 3.42E-06 |
| RIMS4 | 49.29713 | -1.93492 | 0.527841 | -3.66572 | 0.000247 | 0.041453 |
| PLAT | 157.5183 | -1.88416 | 0.413248 | -4.5594 | 5.13E-06 | 0.002997 |
| MXRA5 | 673.7183 | -1.87979 | 0.240384 | -7.81997 | 5.28E-15 | 6.48E-11 |
| PCDH11X | 77.17939 | -1.63444 | 0.393766 | -4.15079 | 3.31E-05 | 0.011292 |
| TOX | 59.35325 | -1.56272 | 0.402633 | -3.88125 | 0.000104 | 0.024984 |
| SCUBE3 | 415.2928 | -1.56219 | 0.26989 | -5.78824 | 7.11E-09 | 1.45E-05 |
| NPNT | 105.4931 | -1.55854 | 0.385924 | -4.03845 | 5.38E-05 | 0.015003 |
| ITPR2 | 192.8205 | -1.53241 | 0.292991 | -5.23025 | 1.69E-07 | 0.000208 |
| PMP22 | 110.8797 | -1.47179 | 0.400893 | -3.67128 | 0.000241 | 0.041124 |
| SMOC1 | 264.6917 | -1.44675 | 0.367679 | -3.93481 | 8.33E-05 | 0.020976 |
| KCNJ2 | 93.47651 | -1.42597 | 0.338871 | -4.20802 | 2.58E-05 | 0.009578 |
| LRP1B | 408.2599 | -1.42207 | 0.253227 | -5.61581 | 1.96E-08 | 3.00E-05 |
| BNC1 | 134.4849 | -1.38586 | 0.321224 | -4.3143 | 1.60E-05 | 0.007015 |
| CCNG2 | 166.7997 | -1.37201 | 0.290158 | -4.7285 | 2.26E-06 | 0.001542 |
| NAALAD2 | 92.52872 | -1.32826 | 0.353884 | -3.75336 | 0.000174 | 0.033448 |
| EGR1 | 989.0697 | -1.26353 | 0.27016 | -4.67697 | 2.91E-06 | 0.00188 |
| UNC5C | 679.7226 | -1.254 | 0.31666 | -3.96007 | 7.49E-05 | 0.019984 |
| PCDH9 | 191.3737 | -1.22727 | 0.299522 | -4.09743 | 4.18E-05 | 0.013488 |
| NTRK2 | 140.2454 | -1.22135 | 0.302046 | -4.04358 | 5.26E-05 | 0.015003 |
| ARHGAP28 | 215.8284 | -1.12438 | 0.284573 | -3.9511 | 7.78E-05 | 0.020307 |
| DAB2 | 156.0343 | -1.10424 | 0.287592 | -3.83961 | 0.000123 | 0.027776 |
| KALRN | 231.5837 | -1.07302 | 0.295672 | -3.62907 | 0.000284 | 0.044741 |
| COL6A3 | 810.9889 | -1.01065 | 0.241038 | -4.1929 | 2.75E-05 | 0.009734 |
| MKI67 | 474.8936 | -0.91437 | 0.245653 | -3.7222 | 0.000197 | 0.035634 |
| HMCN1 | 511.4645 | -0.90512 | 0.247655 | -3.65475 | 0.000257 | 0.042682 |
| TCF4 | 393.814 | -0.90377 | 0.231775 | -3.89932 | 9.65E-05 | 0.02367 |
| SESN3 | 951.6673 | -0.90163 | 0.196144 | -4.59677 | 4.29E-06 | 0.002632 |
| BMP4 | 388.7607 | -0.83077 | 0.223954 | -3.70954 | 0.000208 | 0.036921 |
| FREM2 | 3094.447 | -0.72884 | 0.199808 | -3.6477 | 0.000265 | 0.043285 |
| ATP13A3 | 1336.835 | 0.761789 | 0.211385 | 3.603797 | 0.000314 | 0.046922 |
| SPEG | 1087.916 | 0.813882 | 0.214508 | 3.794177 | 0.000148 | 0.030804 |
| ADCY6 | 739.9975 | 0.839866 | 0.213525 | 3.933338 | 8.38E-05 | 0.020976 |
| HCN4 | 1701.113 | 0.917581 | 0.244432 | 3.753937 | 0.000174 | 0.033448 |
| VEGFA | 1417.398 | 0.951794 | 0.216357 | 4.399191 | 1.09E-05 | 0.005332 |
| JPH2 | 557.6006 | 0.954092 | 0.259016 | 3.68352 | 0.00023 | 0.039751 |
| ANKRD1 | 366.0845 | 0.963788 | 0.266646 | 3.614486 | 0.000301 | 0.046673 |
| SMAD9 | 221.9178 | 0.978237 | 0.270861 | 3.611583 | 0.000304 | 0.046673 |
| MYLK3 | 426.487 | 0.978485 | 0.239309 | 4.088789 | 4.34E-05 | 0.013642 |
| FAM162A | 254.492 | 1.011916 | 0.27838 | 3.635021 | 0.000278 | 0.044289 |
| SNTA1 | 532.6621 | 1.031993 | 0.268954 | 3.83706 | 0.000125 | 0.027776 |
| DES | 379.5385 | 1.038261 | 0.233742 | 4.441908 | 8.92E-06 | 0.004756 |
| FMOD | 141.8295 | 1.088624 | 0.298984 | 3.641082 | 0.000271 | 0.043828 |
| **LBH** | 749.6885 | 1.095298 | 0.24731 | 4.428852 | 9.47E-06 | 0.004843 |
| **PLN** | 255.9963 | 1.152566 | 0.268883 | 4.2865 | 1.82E-05 | 0.007679 |
| **OTOGL** | 92.12968 | 1.233416 | 0.34205 | 3.605953 | 0.000311 | 0.046922 |
| **ILDR2** | 215.5944 | 1.243889 | 0.30089 | 4.134035 | 3.56E-05 | 0.01182 |
| **TMOD1** | 445.9248 | 1.244511 | 0.332684 | 3.740815 | 0.000183 | 0.034622 |
| **TMEM200B** | 153.5161 | 1.246078 | 0.313311 | 3.977134 | 6.98E-05 | 0.019017 |
| **SYNPO2L** | 299.1154 | 1.280779 | 0.29193 | 4.387284 | 1.15E-05 | 0.005416 |
| **B4GALNT3** | 211.4983 | 1.28182 | 0.299736 | 4.276495 | 1.90E-05 | 0.007765 |
| **PTK2B** | 119.289 | 1.291179 | 0.336105 | 3.8416 | 0.000122 | 0.027776 |
| **ADPRHL1** | 606.0543 | 1.309091 | 0.343571 | 3.810251 | 0.000139 | 0.029366 |
| **KEL** | 74.8577 | 1.392672 | 0.368189 | 3.782498 | 0.000155 | 0.031749 |
| **MYL7** | 3505.058 | 1.444363 | 0.266058 | 5.428745 | 5.68E-08 | 7.74E-05 |
| **AHNAK2** | 238.2168 | 1.467382 | 0.409113 | 3.58674 | 0.000335 | 0.049496 |
| **FABP3** | 214.3873 | 1.472752 | 0.361906 | 4.069431 | 4.71E-05 | 0.013767 |
| **AFP** | 1166.336 | 1.514903 | 0.23323 | 6.495332 | 8.29E-11 | 5.08E-07 |
| **APOB** | 672.4562 | 1.582256 | 0.259719 | 6.09219 | 1.11E-09 | 3.42E-06 |
| **GIPR** | 92.75695 | 1.584166 | 0.428182 | 3.699747 | 0.000216 | 0.037826 |
| **MASP1** | 121.8525 | 1.591811 | 0.317228 | 5.017878 | 5.22E-07 | 0.000506 |
| **SERPINA1** | 182.51 | 1.608073 | 0.383694 | 4.191027 | 2.78E-05 | 0.009734 |
| **SPTB** | 99.34901 | 1.67513 | 0.437452 | 3.829292 | 0.000129 | 0.028156 |
| **MB** | 147.0235 | 1.950134 | 0.436063 | 4.47214 | 7.74E-06 | 0.004319 |
| **ACTN3** | 82.63676 | 1.989719 | 0.513253 | 3.876686 | 0.000106 | 0.024984 |
| **PLCXD3** | 109.8085 | 2.05051 | 0.481794 | 4.255992 | 2.08E-05 | 0.008237 |
| **CSRP3** | 99.98391 | 2.092456 | 0.441532 | 4.739081 | 2.15E-06 | 0.001542 |
| **DUOX2** | 48.53778 | 2.813816 | 0.553697 | 5.081865 | 3.74E-07 | 0.000417 |
| **SYNGR3** | 20.01526 | 3.630709 | 0.952305 | 3.81255 | 0.000138 | 0.029366 |
| **ABRA** | 20.21581 | 3.815949 | 0.937148 | 4.071873 | 4.66E-05 | 0.013767 |

**Supplementary Table S4.** : Ventricular ET-1 treatment over control, differential genes

|  | baseMean | log2FoldChange | lfcSE | stat | p-value | p-adjusted |
| --- | --- | --- | --- | --- | --- | --- |
| RBPJL | 0.856158 | -23.6315 | 3.814935 | -6.19447 | 5.85E-10 | 3.04E-06 |
| ADH1A | 0.530105 | -17.887 | 4.025442 | -4.44348 | 8.85E-06 | 0.005523 |
| ISL2 | 1.109428 | -17.6389 | 3.471295 | -5.08137 | 3.75E-07 | 0.000487 |
| DIPK2B | 16.22717 | -3.02832 | 0.785779 | -3.8539 | 0.000116 | 0.032268 |
| GRID1 | 28.71 | -2.66177 | 0.586665 | -4.53713 | 5.70E-06 | 0.003706 |
| PTGER3 | 36.48847 | -2.05783 | 0.492551 | -4.17791 | 2.94E-05 | 0.013907 |
| LINC02082 | 52.96861 | -1.91433 | 0.463127 | -4.13348 | 3.57E-05 | 0.015482 |
| PTCHD1 | 76.83048 | -1.8099 | 0.379897 | -4.76419 | 1.90E-06 | 0.001557 |
| EGFEM1P | 195.3621 | -1.76216 | 0.357452 | -4.92979 | 8.23E-07 | 0.000794 |
| KCNH7 | 502.7944 | -1.64494 | 0.224987 | -7.31128 | 2.65E-13 | 4.13E-09 |
| IRX4 | 208.7098 | -1.62404 | 0.308269 | -5.26826 | 1.38E-07 | 0.000269 |
| SPOCK1 | 64.35338 | -1.62265 | 0.409753 | -3.96007 | 7.49E-05 | 0.023447 |
| PLXNA4 | 638.4861 | -1.56988 | 0.306062 | -5.12928 | 2.91E-07 | 0.000481 |
| SORCS3 | 127.6226 | -1.4373 | 0.368053 | -3.90514 | 9.42E-05 | 0.027203 |
| HPSE2 | 122.0173 | -1.42581 | 0.377775 | -3.77423 | 0.000161 | 0.03912 |
| CD93 | 162.917 | -1.35131 | 0.296782 | -4.5532 | 5.28E-06 | 0.003706 |
| KCTD12 | 2567.873 | -1.22496 | 0.178673 | -6.85588 | 7.09E-12 | 5.53E-08 |
| TRH | 321.0296 | -1.16292 | 0.228502 | -5.08932 | 3.59E-07 | 0.000487 |
| BEX1 | 225.175 | -1.15576 | 0.283002 | -4.08393 | 4.43E-05 | 0.018177 |
| CDH5 | 173.191 | -1.14643 | 0.283706 | -4.04089 | 5.32E-05 | 0.020765 |
| HCN1 | 221.1742 | -1.13375 | 0.286348 | -3.95934 | 7.52E-05 | 0.023447 |
| LAMA2 | 1182.987 | -1.12195 | 0.212549 | -5.27854 | 1.30E-07 | 0.000269 |
| MATN2 | 522.8378 | -1.11318 | 0.236808 | -4.70075 | 2.59E-06 | 0.002022 |
| CPT1A | 137.8244 | -1.10056 | 0.277369 | -3.96786 | 7.25E-05 | 0.023447 |
| FREM1 | 455.7395 | -1.09533 | 0.247362 | -4.42806 | 9.51E-06 | 0.005705 |
| LRRC10 | 197.517 | -1.08499 | 0.252205 | -4.30203 | 1.69E-05 | 0.009104 |
| ADGRG2 | 213.2524 | -1.08203 | 0.281015 | -3.85043 | 0.000118 | 0.032268 |
| NREP | 2801.726 | -1.03845 | 0.192794 | -5.3863 | 7.19E-08 | 0.000187 |
| BMP5 | 1348.577 | -1.00881 | 0.242992 | -4.15163 | 3.30E-05 | 0.015145 |
| VCAM1 | 1000.819 | -1.00011 | 0.267701 | -3.73594 | 0.000187 | 0.044201 |
| SPHKAP | 651.4761 | -0.97122 | 0.213798 | -4.54271 | 5.55E-06 | 0.003706 |
| EFNB3 | 399.3339 | -0.97091 | 0.228907 | -4.24151 | 2.22E-05 | 0.011172 |
| FHOD3 | 1183.769 | -0.94003 | 0.187237 | -5.02052 | 5.15E-07 | 0.000574 |
| RNF207 | 310.7753 | -0.93465 | 0.232046 | -4.02786 | 5.63E-05 | 0.021415 |
| ERBB4 | 1265.213 | -0.89225 | 0.180696 | -4.93785 | 7.90E-07 | 0.000794 |
| CORIN | 492.0912 | -0.891 | 0.215448 | -4.13557 | 3.54E-05 | 0.015482 |
| HCN4 | 1597.482 | -0.88414 | 0.208944 | -4.23146 | 2.32E-05 | 0.011318 |
| FAM189A2 | 948.8557 | -0.86163 | 0.212536 | -4.05405 | 5.03E-05 | 0.020134 |
| PLCL2 | 381.6259 | -0.85539 | 0.224349 | -3.81279 | 0.000137 | 0.035723 |
| LGALS3BP | 421.4418 | -0.84578 | 0.2125 | -3.98015 | 6.89E-05 | 0.023447 |
| PPP1R13B | 777.0512 | -0.82818 | 0.213839 | -3.8729 | 0.000108 | 0.030502 |
| ADGRB2 | 576.4719 | -0.76738 | 0.190808 | -4.02175 | 5.78E-05 | 0.021455 |
| ZNF436 | 407.9811 | -0.76455 | 0.202798 | -3.77002 | 0.000163 | 0.039174 |
| SFRP1 | 1119.771 | -0.75813 | 0.200857 | -3.77447 | 0.00016 | 0.03912 |
| ATP1B1 | 1052.845 | -0.74929 | 0.172277 | -4.34935 | 1.37E-05 | 0.007889 |
| MTUS2 | 600.5205 | -0.73122 | 0.196606 | -3.7192 | 0.0002 | 0.04653 |
| CACNA1C | 1163.62 | -0.70389 | 0.183887 | -3.82784 | 0.000129 | 0.034178 |
| KIF26B | 2514.659 | -0.69355 | 0.1744 | -3.97676 | 6.99E-05 | 0.023447 |
| VCAN | 14055.85 | -0.6879 | 0.166709 | -4.12637 | 3.69E-05 | 0.015537 |
| CNTNAP2 | 238.6141 | 1.086379 | 0.255363 | 4.254259 | 2.10E-05 | 0.010906 |
| KIF5C | 192.5288 | 1.190366 | 0.303686 | 3.919729 | 8.86E-05 | 0.026584 |
| GREB1L | 328.903 | 1.241295 | 0.287187 | 4.32225 | 1.54E-05 | 0.008604 |
| A2M | 539.0483 | 1.258478 | 0.259666 | 4.846521 | 1.26E-06 | 0.001089 |
| AIF1L | 142.3834 | 1.262723 | 0.270001 | 4.676739 | 2.91E-06 | 0.002165 |
| ARHGAP11A | 63.22785 | 1.44443 | 0.376744 | 3.833982 | 0.000126 | 0.03391 |
| IRX2 | 105.9969 | 1.731042 | 0.35183 | 4.920112 | 8.65E-07 | 0.000794 |
| **LIN28A** | 558.9515 | 1.897905 | 0.320323 | 5.924966 | 3.12E-09 | 1.18E-05 |
| **NUF2** | 32.51696 | 2.202695 | 0.550282 | 4.002846 | 6.26E-05 | 0.022704 |
| **PTPRZ1** | 88.31804 | 2.235186 | 0.443527 | 5.039569 | 4.67E-07 | 0.00056 |
| **OTX1** | 62.14143 | 2.308019 | 0.579208 | 3.984782 | 6.75E-05 | 0.023447 |
| **ALPL** | 31.68836 | 2.375996 | 0.629503 | 3.774399 | 0.00016 | 0.03912 |
| **PLP1** | 44.61831 | 4.547992 | 0.771786 | 5.892815 | 3.80E-09 | 1.18E-05 |
| **ELAPOR1** | 13.05028 | 4.636459 | 1.219167 | 3.802974 | 0.000143 | 0.03656 |
| **DMBX1** | 25.48825 | 5.1165 | 0.999666 | 5.118207 | 3.08E-07 | 0.000481 |
| **FABP7** | 10.84472 | 5.756118 | 1.47019 | 3.91522 | 9.03E-05 | 0.026584 |
| **MAD2L1P1** | 3.078773 | 15.77035 | 4.013118 | 3.929699 | 8.51E-05 | 0.026014 |
| **SLC36A2** | 0.492018 | 15.98102 | 4.024342 | 3.97109 | 7.15E-05 | 0.023447 |

## Supplementary Videos - legends

**Video S1** – Bright field video of atrial EBs cultured with ET-1 treatment from day 4 to day 7.

**Video S2** - Bright field video of ventricular EBs cultured with ET-1 treatment from day 4 to day 7.

**Video S3** -Wavefront propagation of the depolarization in control atrial EBs, measured with the voltage sensitive FluoVolt dye at differentiation day 14.

**Video S4** - Wavefront propagation of the depolarization in ET-1 treated atrial EBs (from day 4 to 7), measured with the voltage sensitive FluoVolt dye at differentiation day 14.

**Video S5** - Wavefront propagation of the depolarization in control ventricular EBs, measured with the voltage sensitive FluoVolt dye at differentiation day 14.

**Video S6** - Wavefront propagation of the depolarization in ET-1 treated ventricular EBs (from day 4 to 7), measured with the voltage sensitive FluoVolt dye at differentiation day 14.

